# Interspecific divergence of circadian properties in duckweed plants

**DOI:** 10.1101/2021.10.19.465055

**Authors:** Minako Isoda, Shogo Ito, Tokitaka Oyama

## Abstract

- The circadian clock system is widely conserved in plants; however, divergence in circadian rhythm properties is poorly understood. We conducted a comparative analysis of the circadian properties of closely related duckweed species.
- Using a particle bombardment method, a circadian bioluminescent reporter was introduced into duckweed plants. We measured bioluminescence circadian rhythms of eight species of the genus *Lemna* and seven species of the genus *Wolffiella* at various temperatures (20, 25, and 30 °C) and light conditions (constant light or constant dark). *Wolffiella* species inhabit relatively warm areas and lack some tissues/organs found in *Lemna* species.
- *Lemna* species tended to show robust bioluminescence circadian rhythms under all conditions, while *Wolffiella* species showed lower rhythm stability, especially at higher temperatures. For *Lemna*, two species (*L. valdiviana* and *L. minuta*) forming a clade showed relatively lower circadian stability. For *Wolffiella*, two species (*W. hyalina* and *W. repanda*) forming a clade showed extremely long period lengths.
- The circadian properties of species primarily reflect their phylogenetic positions. The relationships between geographical and morphological factors and circadian properties are also suggested.

## Introduction

Circadian clocks are endogenous timekeeping systems that allow organisms to anticipate the daily and seasonal changes surrounding them. Many organisms, from cyanobacteria to humans, have a circadian clock with a period of approximately 24 h, and the circadian clock modulates the timing of various physiological phenomena. The circadian clock of a plant under day-night cycles is synchronized to diurnal changes in external information, such as light, dark, and temperature, and this regulates many physiological processes including leaf movement, stomatal opening and closing, and petal opening (Inoue *et al*., 2018). In *Arabidopsis thaliana*, more than 20 clock-related components have been identified (Hsu & Harmer, 2014). The core circadian clock genes, including *CIRCADIAN CLOCK ASSOCIATED 1* (*CCA1*) and *PSEUDO-RESPONSE REGULATOR* (*PRR*) family genes, *GIGANTEA* (*GI*), *LUXARRHYTHMO* (*LUX*), *EARLY FLOWERING 3* (*ELF3*), and *EARLY FLOWERING 4* (*ELF4*), form a regulatory network with transcription-translation feedback loops (Pokhilko *et al*., 2012; Nohales & Kay, 2016; Sanchez *et al*., 2020). These genes have been identified in species as diverse as green algae (*Ostreococcus tauri*), charophytes, moss (*Physcomitrella patens*), and many higher plants (Song *et al*., 2010; Linde *et al*., 2017). Indeed, from algae to eudicots, diurnal transcriptomes are remarkably similar despite the large phylogenetic distances and differences in morphological complexity and habitat (Ferrari *et al*., 2019). Thus, the circadian clock mechanisms studied in *Arabidopsis* are widely conserved in the plant kingdom. Despite this, even within the same species, *Arabidopsis thaliana*, a latitudinal cline has been observed in the period which displays a large natural variation (Michael *et al*., 2003). By comparing leaf movements, Müller *et al*. showed that the circadian clock of cultivated tomato species runs more slowly than that of its wild relatives (Müller *et al*., 2016). These results suggest that circadian phenomena display variations important for local adaptation. Plants have evolved to adapt to external environments, resulting in a variety of natural variations (Müller *et al*., 2016; Greenham *et al*., 2017). The light and temperature conditions experienced by plants are largely dependent on local climate, and light and temperature are well-known input signals for plant circadian clocks (Inoue *et al*., 2018). While natural variations in plant circadian properties have been analyzed for strains/accessions distributed across relatively similar climatic environments, this has not been done for species distributed across regions with dramatically different climates.

Lemnaceae, commonly known as duckweed, is a group of small aquatic monocotyledonous plants widely distributed in tropical, arid, temperate, and subarctic areas. Duckweed includes 36 species representing the five genera *Spirodela, Landoltia, Lemna, Wolffiella*, and *Wolffia* (Sree *et al*., 2016; Bog *et al*., 2020). Morphologically, duckweed is traditionally classified into two groups—Lemnoideae, which includes the genera *Spirodela, Landoltia*, and *Lemna*; and Wolffioideae, which includes the genera *Wolffiella* and *Wolffia* (Landolt, 1986). Lemnoideae species have nerves (leaf veins) and roots and two budding pouches in a frond (leaf-like structure). In contrast, Wolffioideae species lack nerves and roots and have one budding pouch in a frond (Les *et al*., 1997). Lemnoideae species are widely distributed from low to high latitudes, whereas Wolffioideae species are mainly found in low-latitude areas. Duckweed has been used for physiological studies since the 1950s because of its size and rapid growth (Acosta *et al*., 2021). With respect to circadian rhythm studies, *Lemna* plants have been used for the analysis of physiological rhythms and molecular studies on the circadian clock machinery (Miyata & Yamamoto, 1969; Hillman, 1970; Kondo & Tsudzuki, 1978). Clock-related genes, including *LHY*s, *PRR*s, *GIGANTIA* (*GI*), and *EARLY FLOWERING3* (*ELF3*), have been identified in several duckweed plants (Miwa *et al*., 2006; Michael *et al*., 2021). To observe circadian rhythms in duckweed, bioluminescence monitoring systems have been used (Miwa *et al*., 2006; Serikawa *et al*., 2008). A bioluminescence reporter, in which the firefly luciferase gene was driven by the *Arabidopsis CCA1* promoter (*AtCCA1::LUC+*), was introduced into duckweed using particle bombardment, and the bioluminescence of the plants grown on a luciferin-containing medium was automatically monitored for a week or more (Muranaka & Oyama, 2016). Using the bioluminescence monitoring system, a comparative analysis of circadian rhythms in nine strains of duckweed representing five species and four genera (*Spirodela, Landoltia, Lemna*, and *Wolffia*) was performed (Muranaka *et al*., 2015). Based on the results of this previous study, under light/dark conditions, *AtCCA1::LUC+* reporter activity showed diurnal rhythms peaking in the early morning in each strain. Under constant light conditions, *AtCCA1::LUC+* reporter activity showed more variable rhythmic behavior, specifically robust/mildly-dampened rhythms in strains of *Spirodela polyrhiza, Landoltia punctata*, and *Lemna gibba*, and the Nd strain of *Lemna aequinoctialis*; unstable rhythms in *Wolffia columbiana*; and severely dampened rhythms in the 6746 strain of *Lemna aequinoctialis*. Thus, even within the same species, different circadian behaviors are observed. Furthermore, under constant dark conditions, *AtCCA1::LUC+* reporter activity showed dampened circadian rhythms in all strains. The period length was also longer under constant dark conditions than under constant light conditions in all strains except *Wolffia columbiana*. These results suggest that circadian properties, such as period length and stability, widely vary among duckweed plants and may be optimized for specific environmental conditions. Interestingly, *Wolffia australiana* has been reported to carry approximately half the numbers of light-signaling and circadian clock genes than *Spirodela polyrhiza* (Michael *et al*., 2021). Circadian systems of *Wolffia* species may have dramatically diversified when the genus arose. Taken together, interspecific divergence found in the circadian properties of duckweed plants seems to occur at different phylogenetic levels. It is not yet known, however, how these circadian properties are geographically related between different duckweed habitats.

In this study, we aimed to capture the interspecific divergence in the circadian properties of duckweed plants. We focused on two genera, *Lemna* (Lemnoideae) and *Wolffiella* (Wolffioideae). Among the Lemnoideae, the genus *Spirodela* has two species and the genus *Landoltia* has only one species; the *Lemna* genus contains 12 species, which makes it highly suitable for comparing interspecific divergence in circadian properties (Acosta *et al*., 2021). In Wolffioideae, the sizes of *Wolffiella* plants are larger than those of *Wolffia*, which are too small to handle experimentally. It has been reported that the habitats of these genera overlap in various combinations (Landolt, 1986). Species in each genus were selected to cover their full distributional ranges. We characterized circadian rhythms in eight *Lemna* species and seven *Wolffiella* species by monitoring bioluminescence circadian reporters under constant and light/dark-entrainment conditions at different temperatures (20, 25, and 30 °C). We show that while every species has the potential for self-sustained oscillation and entrainability, the stability of circadian rhythms was highly dependent on phylogeny. Based on our results, we suggest that the circadian properties of *Lemnaceae* are phylogenetically restricted rather than geographically restricted.

## Materials and Methods

### Plant materials and growth conditions

*Wolffiella lingulata* 7547 and *W. oblonga* 7201 were provided by the Rutgers Duckweed Stock Cooperative (http://www.ruduckweed.org/); *W. caudata* 9139, *W. neotropica* 7279, *W. welwitschii* 7644, *W. hyalina* 9525, *W. repanda* 9122, *Lemna obscura* 8892, *L. turionifera* 6619, *L. japonica* 8695, *L. disperma* 7767, *L. valdiviana* 9475, and *L. minuta* 9476 strains were provided by the Landolt Duckweed Collection (Dr. Walter Lämmler, http://www.duckweed.ch/); *L. minor* 5512 was provided by Dr. Masaaki Morikawa (Hokkaido University). The *L. gibba* p8L strain is a pure line (eight generations of selfing) produced from the *L. gibba* 7741 (G3) strain (Muranaka *et al*., 2015). All plants were aseptically kept on modified NF medium with 1% sucrose and 5 mM MES [2-(N-morpholino)-ethanesulfonic acid] as previously described for *L. gibba* (Muranaka *et al*., 2015). Plants were cultured in 8 ml of NF medium with continuous light at 25 ± 1 °C, with light supplied by fluorescent lamps (FLR40SEXW/M/36-HG; NEC) at approximately 50 µmol m^−2^ s^−1^.

In the growth experiment, the plants were precultured in Bio Multi Incubator (LH-30-8CT; NK system) under a 12 h light/12 h dark cycle for two days. The light intensity was approximately 40 µmol m^−2^ s^−1^ with a fluorescent lamp (FML13EX-N DK10; Hitachi), and the temperature was maintained at either 20, 25, or 30 °C. After preculture, the plants were placed in 12-well plates (one colony per well) with 4 ml of NF medium and grown under constant light conditions at each experimental temperature. After one week, the number of plant colonies in each well was counted.

### Luciferase reporter constructs

We used *pUC-AtCCA1::intron-LUC+:NosT* (*AtCCA1::LUC+*) (Watanabe *et al*., 2021) and *pUC18 pAtCCR2:intronLUC+:NosT* (*AtCCR2::LUC+*) as circadian bioluminescent reporters. The promoter region of the *Arabidopsis CCR2* gene was amplified by PCR with the genomic DNA template (*Arabidopsis thaliana* Col-0) and the primers (5’-TGGATCCACCGTGTGAGTTGGTAGCG-3’ and 5’-AGGCGCGCCTGAAATTTGAAAAGAAGATCTAAG-3’) (Strayer *et al*., 2000). The 1.4-kb DNA fragment was cloned into pENTR 5’-TOPO (Invitrogen) with an additional multi-cloning site. Using the LR reaction, the *AtCCR2* promoter in this plasmid was integrated into pUC18-*intron-LUC+*, in which the *aat*R4-*att*L1 sequence was followed by the *intron-LUC+* and *Nos* terminator (Muranaka *et al*., 2015).

### Particle bombardment experiment and bioluminescence monitoring

Reporter constructs were introduced into the frond cells by particle bombardment, as described previously with minor modifications (Muranaka *et al*., 2015). Briefly, 0.48 mg of gold particles (1.0-μm diameter; Bio-Rad) were coated with 2 μg of plasmid DNA and introduced into plants laid on a 35-mm polystyrene dish using a helium gun device (PDS-1000/He, Bio-Rad) according to the manufacturer’s instructions (vacuum, 27 mmHg; helium pressure, 450 psi). After particle bombardment, the 35-mm dish was filled with 4 ml of modified NF medium containing D-luciferin (0.1 mM potassium salt, Wako) and set in a bioluminescence monitoring system in an incubator (KCLP-1000I-CT, NK system). Bioluminescence monitoring was performed as described previously (Muranaka *et al*., 2015). The light intensity in the incubator was approximately 30 μmol m^-2^ s ^-1^ and the temperature was maintained at 20 ± 1 °C, 25 ± 1 °C, and 30 ± 1 °C, respectively.

### Estimation of period lengths

A time-series analysis was performed using R 3.4.1 (http://www.R-project.org/). The period lengths under constant conditions were estimated by the fast Fourier transform non-linear least squares method, using 72-h of data (RAE < 0.2) obtained 24 h after changing from light/dark to constant light or constant dark (Muranaka & Oyama, 2016).

## Results

### Phylogenetic and geographic relationships of 15 species of Lemnaceae

We used duckweed species mainly from germplasm collections (see Materials and Methods), selecting seven out of ten *Wolffiella* species and eight out of 12 *Lemna* species (Fig. 1; Fig. S1, S2) (Tippery *et al*., 2015). Phylogenetic studies have shown that *Wolffia* and *Wolffiella* are the most derived genera, and *Spirodela* is the most ancestral genus (Les *et al*., 2002; Tippery *et al*., 2015). In the case of *Wolffiella, W. hyalina* and *W. repanda* were phylogenetically close. In the case of *Lemna, L. valdiviana*, and *L. minuta* were phylogenetically close. We selected *Wolffiella* species, which are distributed in tropical, arid, and temperate areas, and *Lemna* species, which are distributed in tropical, temperate, and subarctic areas (Table 1). Of the strains we examined, *L. minor* 5512 was collected from the coldest area, and *W. caudata* 9139, *W. neotropica* 7279, and *L. valdiviana* 9475 were collected from hot areas; *W. welwitschii* 7644, *W. hyalina* 9525, and *W. repanda* 9122 were collected from hot and semi-arid areas. Specifically, *W. caudata* 9139 and *L. valdiviana* 9475 were collected from the same area of Brazil; and *W. lingulata* 7547 and *L. turionifera* 6619 were collected from the same area of the USA.

**Fig. 1.**
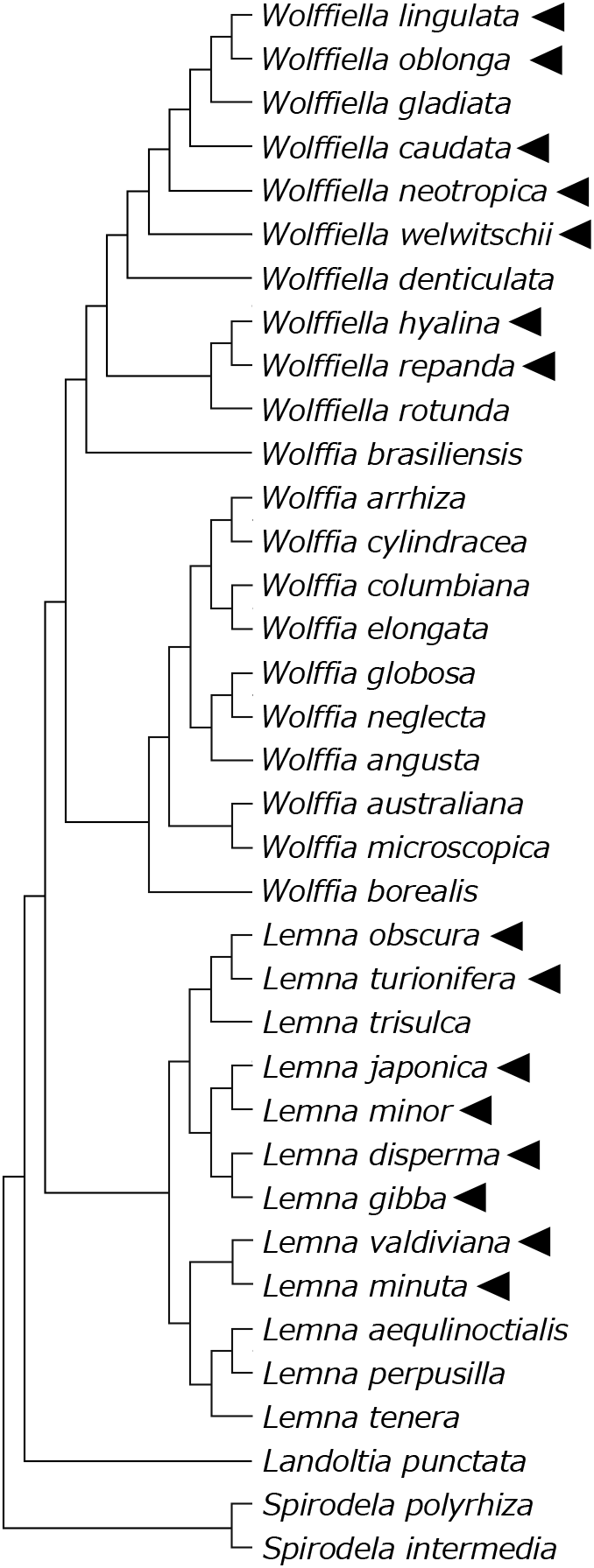
Phylogenetic relationship of the family Lemnaceae. A phylogenetic tree with every species of duckweed (Sree *et al*., 2016, Bog *et al*., 2020) based on Les *et al*. (2002) and Tippery *et al*. (2015). Arrowheads indicate the species used in this study.

**Table 1.**
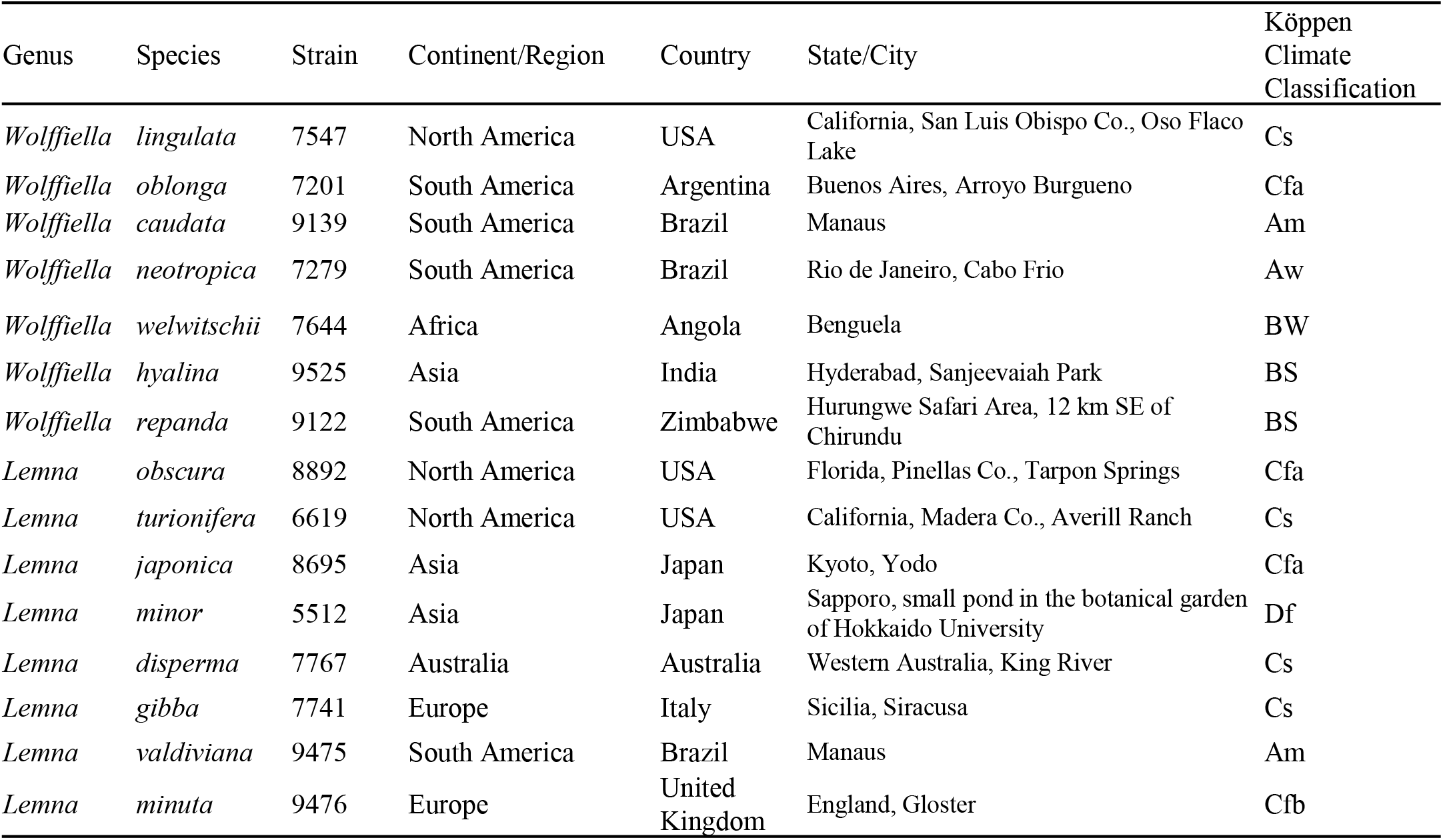
Geographical location and Köppen climate classification on the species used in this study.

### Stability of the circadian rhythms of *Wolffiella* species under constant conditions at different temperatures

Duckweed plants that were cultured under constant light conditions at 25 °C (a standard temperature for duckweed culture) were used for the gene introduction experiments. The *AtCCA1::LUC+* reporter was introduced into the plants by particle bombardment (Muranaka *et al*., 2013). The plants were then placed into an automatic monitoring system set in a growth chamber at 20, 25, or 30 °C. During the monitoring period, these plants were entrained to 12 h dark/ 12 h light cycles and then released into constant light or constant dark conditions (Fig. 2a). We monitored the reporter activities of *AtCCA1::LUC+* in the seven strains of *Wolffiella* (Fig. 2). At 25 °C, every species showed rhythmicity under both constant light and constant dark conditions (Fig. 2b–h). At 30 °C, except for *W. hyalina*, all *Wolffiella* species showed dampened bioluminescence rhythms or arrhythmic reporter activity under constant light conditions (Fig. 2b–h). Here, dampened rhythms indicate bioluminescence traces whose amplitude decreases more rapidly than the degree of alteration in luminescence level. These bioluminescence traces without any clear peaks (excluding the peak around light-on) in the first 48 h under constant conditions are defined as arrhythmic reporter activity (Table S1). At 30 °C, arrhythmic reporter activity or dampened rhythms in these *Wolffiella* species were also observed in the bioluminescence monitoring experiments using the evening-expressed reporter *AtCCR2::LUC*, whereas the activity of this reporter showed rhythmicity at 20 and 25 °C (Fig. S3). These results strongly suggest that the stability of circadian rhythms is lost or severely decreased at higher temperatures, irrespective of circadian reporters.

**Fig. 2.**
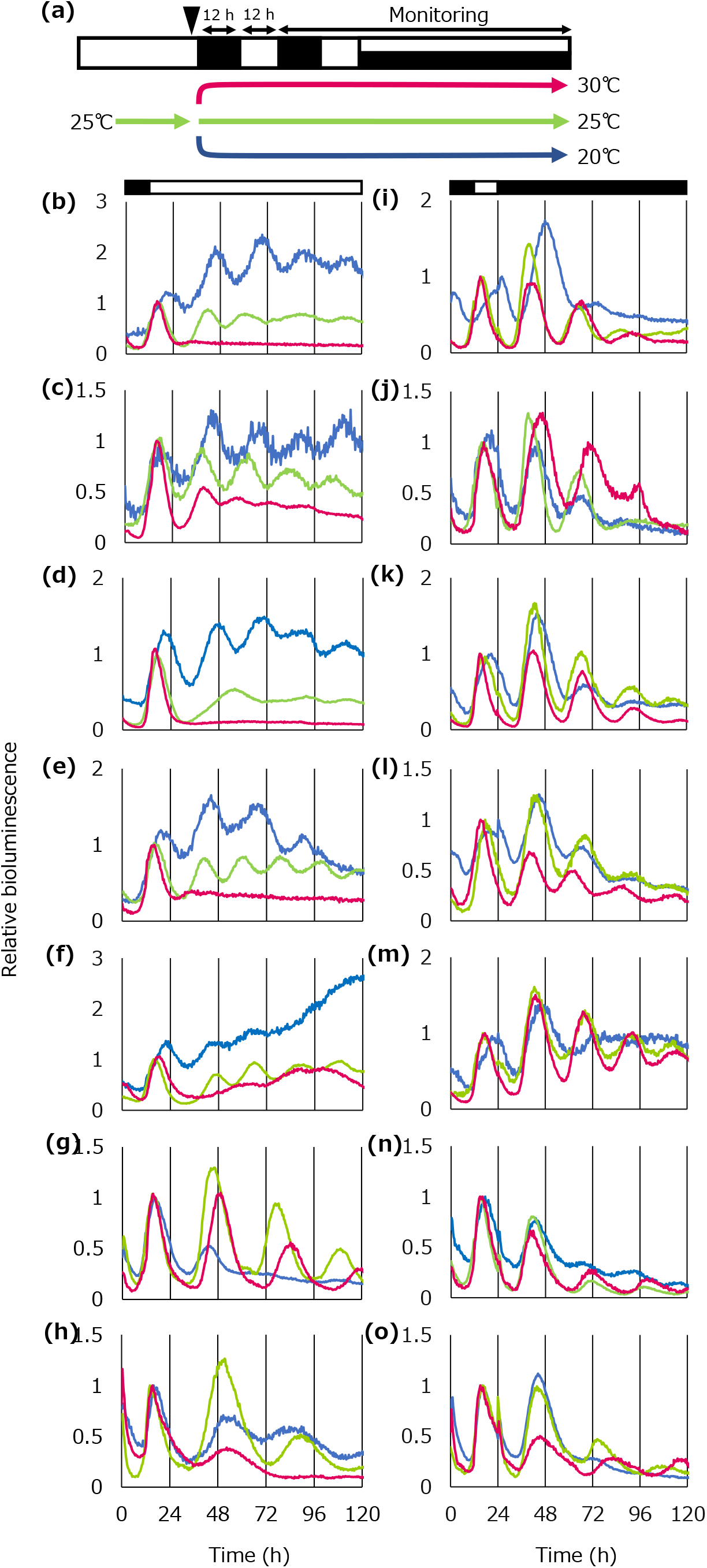
Bioluminescence of *AtCCA1::LUC+* under constant light (b–h) and constant dark (i–o) conditions at various temperatures. (a) Scheme of experimental methods for bioluminescence monitoring. Plants cultured in NF medium under constant light conditions were subjected to gene transfection and then entrained to two 12-h dark/12-h light cycles and released into constant light or constant dark conditions. White and black bars indicate light and dark periods, respectively. *AtCCA1::LUC+* was introduced into *W. lingulata* (b, i), *W. oblonga* (c, j), *W. caudata* (d, k), *W. neotropica* (e, l), *W. welwitschii* (f, m), *W. hyalina* (g, n), and *W. repanda* (h, o). The red, green, and blue colors indicate 30, 25, and 20 °C, respectively. The representative time-series dataof four replicates in two independent experiments is shown for each species/condition. The data are presented as relative values with the highest value between 12 h and 36 h set to 1.

The plants of all *Wolffiella* species grew faster at 30 °C than at lower temperatures (Fig. S4). Although this higher temperature impaired circadian rhythmicity, it was suitable for the growth of *Wolffiella* species. Indeed, the daily maximum temperatures in the natural habitats of this genus exceed 28 °C for more than three months of the year (Landolt, 1986). In contrast, all the *Wolffiella* species showed rhythmicity at 30 °C under constant dark conditions, suggesting that the destabilization of circadian rhythms was dependent on light (Fig. 2i–o). In the entrainment 12-h light/12-h dark conditions, every species showed a clear diurnal rhythm at 30 °C (Fig. S5). At 20 °C, *W. neotropica, W. welwitschii, W. hyalina*, and *W. repanda* showed dampened rhythms under both constant light and constant dark conditions (Fig. 2e–h, l–o). *W. lingulata, W. oblinga*, and *W. caudata* showed circadian rhythms under constant light conditions, while they showed dampened rhythms under constant dark conditions (Fig. 2b–d, i–k). These results indicate that the stability of *Wolffiella* circadian rhythms is highly dependent on light and temperature, and the magnitude of their effects varies among species.

### Stability of the circadian rhythms of *Lemna* species in constant conditions at different temperatures

We monitored the reporter activity of *AtCCA1::LUC+* in the eight *Lemna* strains (Fig. 3). Every *Lemna* species showed bioluminescence rhythms, including dampened rhythms under both constant light and dark conditions at all temperatures. At 25 °C, only *L. turionifera* showed dampened rhythms when monitored under constant dark conditions.

**Fig. 3.**
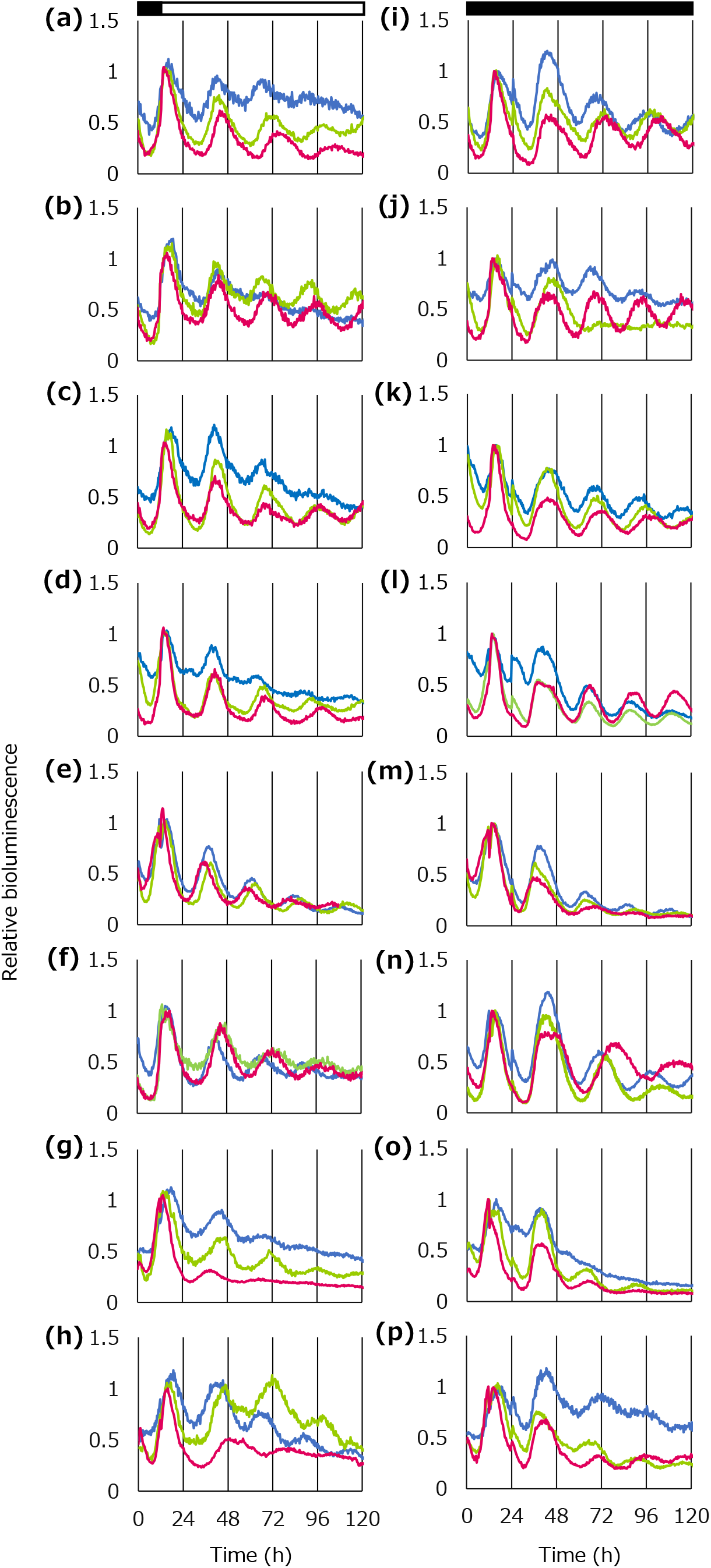
Bioluminescence of *AtCCA1::LUC+* under constant light (a–h) and constant dark (i–p) conditions at various temperatures. *AtCCA1::LUC+* was introduced into (a, i) *L. obscura*, (b, j) *L. turionifera*, (c, k) *L. japonica*, (d, l) *L. minor*, (e, m) *L. disperma*, (f, n) *L. giabba*, (g, o) *L. valdiviana*, and (h, p) *L. minuta*. The red, green, and blue colors indicate 30, 25, and 20 °C, respectively. Plants cultured in NF medium under constant light conditions were subjected to gene transfection and then entrained to two 12-h dark/12-h light cycles and released into constant light or constant dark conditions; white and black bars indicate light and dark periods, respectively. The representative timeseries data of four replicates in two independent experiments are shown for each species/condition. The data are presented as relative values with the highest value between 12 h and 36 h set to 1.

At 30 °C, *L. valdiviana* showed dampened rhythms under both constant light and dark conditions, and *L. minuta* showed dampened rhythms under constant light conditions (Fig. 3g, h, o). At 20 °C, *L. disperma, L. gibba*, and *L. minuta* showed robust rhythms, and the remaining species showed dampened rhythms under constant light conditions (Fig. 3 a– h). In contrast, only *L. valdiviana* and *L. minuta* showed dampened rhythms under constant dark conditions (Fig. 3o, p). In summary, *Lemna* species showed a tendency to maintain robust rhythms at 25 °C or above, while *L. valdiviana* and *L. minuta* showed unstable circadian rhythms at low and high temperatures. The observed high stability in the *Lemna* species under constant light at 30 °C contrasts with the low stability of the *Wolffiella* species.

### Temperature compensation of period lengths in Lemnaceae

We analyzed the period lengths under constant light or dark conditions at the three experimental temperatures (Fig. 4a, b for *Wolffiella*; Fig. 5a, b for *Lemna*). The period lengths of the *Wolffiella* species were strongly affected by light and temperature conditions, showing a wide variation among species (Fig. 3a, b). At 25 °C, under constant light conditions, *W. caudata, W. hyalina*, and *W. repanda* showed extremely long periods (> 30 h), while *W. lingulata, W. oblonga, W. neotropica*, and *W. welwitschii* showed relatively short periods (≤ 24 h) (Fig. 4a, green bars). Under constant dark conditions, the period lengths of *W. lingulata, W. oblonga, W. caudata, W. neotropica*, and *W. welwitschii* were close to 24 h, while those of *W. hyalina* and *W. repanda* were close to 30 h (Fig. 4b, green bars). At 20 °C, the period lengths varied greatly among species (ranging from 19 to 37 h) under constant light conditions. Interestingly, the period lengths of all species were approximately 24 h under constant dark conditions (Fig. 4a, b, blue bars). At 30 °C, under constant light conditions, the period length of *W. hyalina* was estimated to be approximately 36 h and that of *W. oblonga* was approximately 24 h (Fig. 4a, red bars). Under constant dark conditions, the period lengths of *W. lingulata, W. oblonga, W. caudata, W. neotropica*, and *W. welwitschii* were approximately 24 h, while those of *W. hyalina* and *W. repanda* were longer than 30 h (Fig. 4b, red bars). *W. hyalina* and *W. repanda* showed longer period lengths than the other *Wolffiella* species under all conditions. This suggests that with respect to circadian properties, *W. hyalina* and *W. repanda* can be distinguished from other *Wolffiella* species.

**Fig. 4.**
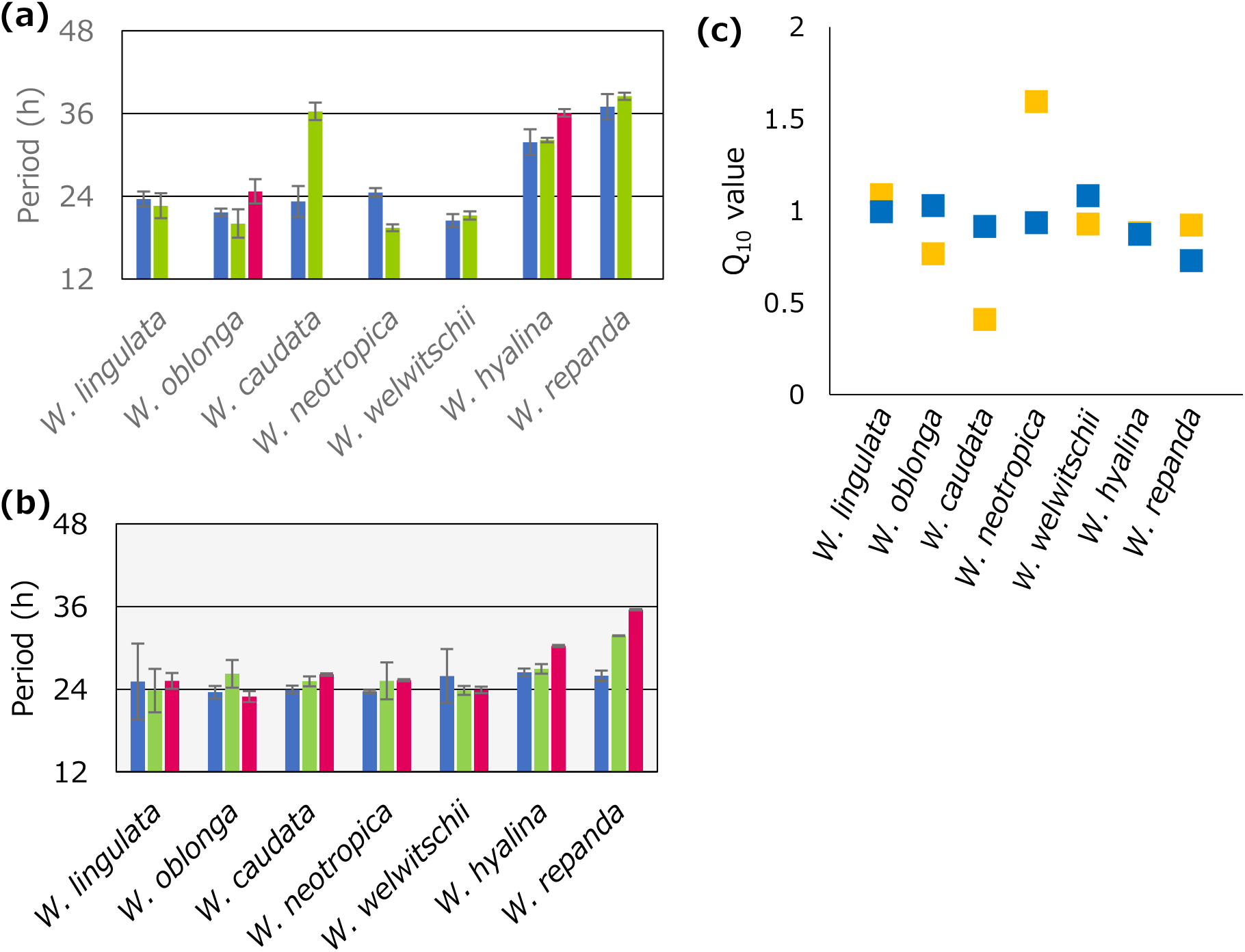
Interspecific divergence of period lengths among *Wolffiella* species. (a) Period lengths under constant light conditions. (b) Period lengths under constant dark conditions. Red, green, and blue colors indicate 30, 25, and 20 °C, respectively. The data represent the mean ± SD (*n* = 4). (c) Q_10_ values of *Wolffiella* species under constant light and constant dark conditions. The yellow and blue squares indicate constant light and constant dark conditions, respectively.

**Fig. 5.**
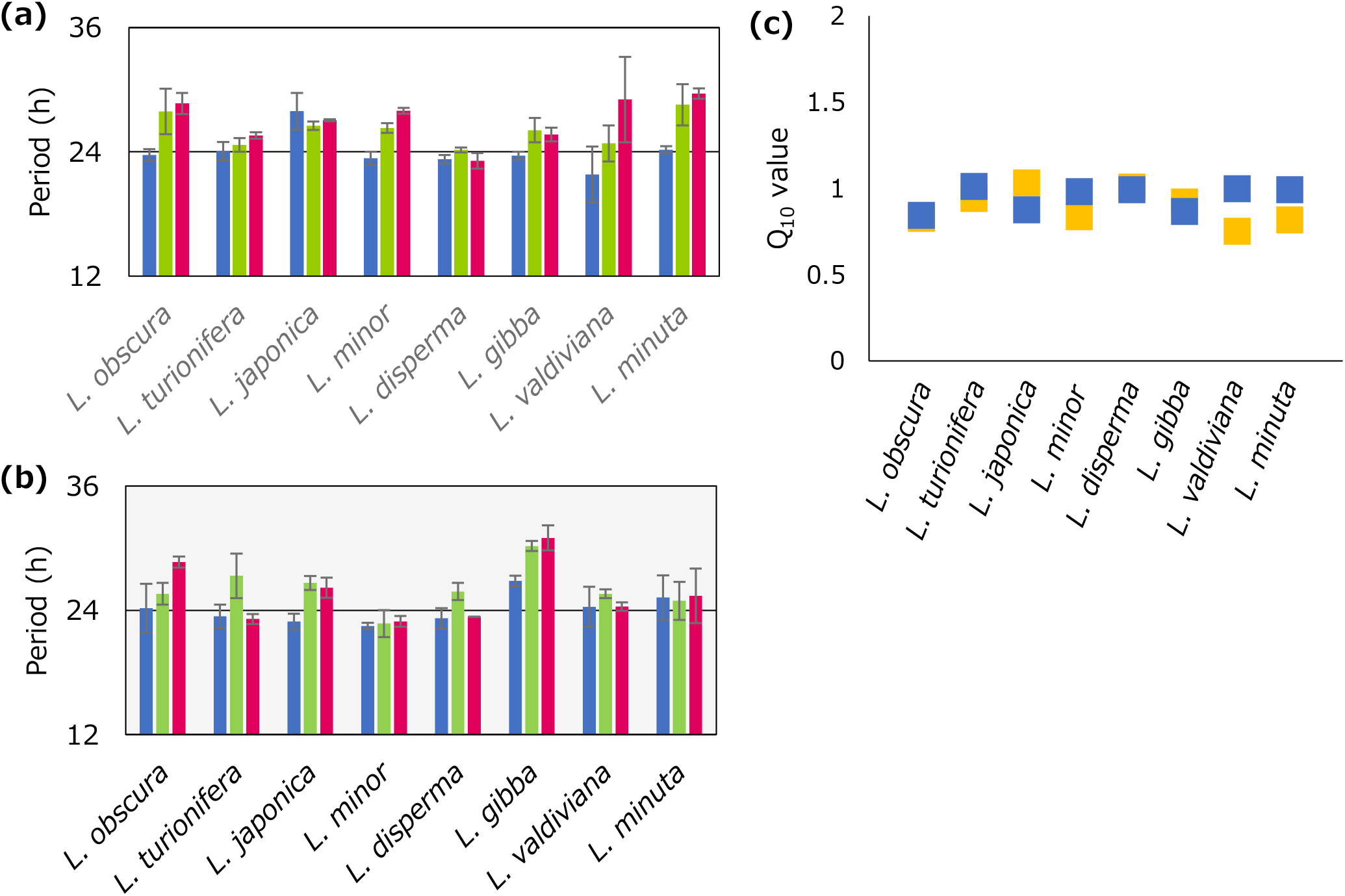
Interspecific divergence of period lengths among *Lemna* species. (a) Period lengths under constant light conditions. (b) Period lengths under constant dark conditions. Red, green, and blue bars indicate 30, 25, and 20 °C, respectively. The data represent the mean ± SD (*n* = 4). (c) Q_10_ values of *Lemna* species under constant light and constant dark conditions. The yellow and blue squares indicate constant light and constant dark conditions, respectively.

The Q_10_ temperature coefficient is the change in the reaction rate with a temperature increase of 10 °C. The Q_10_ value of most biological reactions ranges between 2 and 3, while that of circadian rhythms is close to 1. The Q_10_ values of the *Wolffiella* species under constant light conditions were between 0.40 and 1.59, while those under constant dark conditions were between 0.73 and 1.08 (Fig. 4c). The Q_10_ values of *W. oblonga, W. caudata* and *W. neotropica* were closer to 1 under constant dark conditions than under constant light conditions. This suggests that the temperature compensation of circadian rhythms in *Wolffiella* species is influenced by light conditions.

In the case of the *Lemna* species, the period lengths at 25 °C under constant light conditions were between 24 h and 28 h, and those in constant dark conditions were between 21 h and 30 h (Fig. 5a, b, green bars). In contrast to the *Wolffiella* species, the period lengths among the *Lemna* species varied more under constant dark conditions than under constant light conditions at 25 °C. At 20 °C, the period lengths were approximately 24 h under both constant light and constant dark conditions (Fig. 5a, b, blue bars). At 30 °C, *L. valdiviana* and *L. minuta* showed period lengths close to 30 h under constant light conditions (Fig. 5a, red bars); *L. gibba* also showed period lengths longer than 30 h under constant dark conditions; and the period lengths of the other species were approximately 24 h at 30 °C. The Q_10_ values of the *Lemna* species under constant light conditions ranged between 0.75 and 1.03, while those under constant dark conditions ranged between 0.84 and 1.01 (Fig. 5c). Such interspecific divergence indicates that the temperature compensation of circadian rhythms in *Lemna* is more robust than that of *Wolffiella*, especially under constant light conditions.

### Phylogenetic and geographic comparison of circadian properties

Phylogenetically close species of *Wolffiella* and *Lemna* showed similar circadian properties (Fig. 1; Table S1). In the case of *Wolffiella*, the circadian properties of *W. hyalina* and *W. repanda* were similar in terms of period length and stability. As previously noted, these two species can be distinguished from other species of *Wolffiella* based on their circadian properties, and are phylogenetically close. In the case of *Lemna*, the circadian properties of *L. valdiviana* and *L. minuta* were similar in terms of stability. These two species are phylogenetically close. These results suggest that differences in circadian properties increase with speciation. In particular, the stability of circadian rhythms likely decreased when the *W. hyalina* and *W. repanda* group became distinguished from other *Wolffiella* species.

A geographical factor in the circadian properties can also be broadly identified in *Lemna*; species inhabiting colder climates tend to have more stable circadian rhythms (Table S2). For example, species inhabiting temperate and subarctic areas showed more stable rhythms under high- and low-temperature conditions than those inhabiting the tropics, i.e., *L. valdiviana*. Interestingly, *L. minuta*, which is closest in the phylogenetic tree to *L. valdiviana*, and inhabits temperate areas, also showed unstable rhythms but with a higher level of circadian stability. However, no such tendency was observed in the *Wolffiella* species. These results suggest that interspecific divergence in circadian properties is strongly related to phylogenetic factors rather than geographical factors.

## Discussion

Using a bioluminescence monitoring system, we measured the circadian rhythms of 15 species of *Lemna* and *Wolffiella* under the same strictly controlled conditions. Under various experimental temperatures, we revealed interspecific divergence in period length and stability. *Wolffiella* species tended to show unstable/deviated circadian rhythms, whereas *Lemna* species tended to show stable rhythms. Thus, the circadian properties of the studied species were largely reflected by their phylogenetic position. *Wolffiella* species are morphologically distinct from those of the *Lemna* genus and inhabit relatively low latitude areas, suggesting that differences in circadian properties may further be related to morphological divergence and geographical adaptation.

### Decrease in circadian rhythmicity in *Wolffiella* species

We found highly diverse circadian properties within the *Wolffiella* genus, with most species showing low robustness at high temperature under constant light conditions (Fig. 2). This suggests that mechanisms to maintain circadian rhythmicity under various light/temperature conditions are diversified in *Wolffiella*. Many *Wolffiella* species inhabit low-latitude areas where temperature and day-length variations are small throughout the year (Hut *et al*., 2013). Thus, circadian clocks with high stability may not contribute to the fitness of *Wolffiella* plants, which inhabit relatively constant environments. This may be related to aquatic habitats that commonly experience smaller daily fluctuations than terrestrial environments. Because the circadian rhythms of all *Wolffiella* species were synchronized to the light/dark cycle, even at high temperature (Fig. S5), their unstable circadian clocks must be entrained to day/night cycles in the natural environments.

In addition to the lower stability of circadian rhythms, a wide interspecific divergence in period length was observed in *Wolffiella*. Specifically, under constant light conditions at 25°C, period length varied by approximately 19 h in *Wolffiella* compared to 4 h in *Lemna*. The observed variation between the *Lemna* species is comparable to that reported for other plants; *Arabidopsis thaliana* shows intraspecific variations (natural variation) in period lengths of approximately 7.5 h among a large number of accessions (Michael *et al*., 2003; Edwards *et al*., 2005; Rees *et al*., 2021). *Kalanchoe* also shows interspecific variation in period length of approximately 6 h among six species (Malpas & Jones, 2016). The comparatively large interspecific variation of *Wolffiella* results from the existence of species with extremely long periods, i.e., *W. hyalina* and *W. repanda*, of more than 30 h under constant light conditions(Fig. 4a, b). A very long period has also been reported for *W. hyalina* strain 8640, suggesting that the strain 9525 used in our study has representative circadian properties for this species (Isoda & Oyama, 2018). Interestingly, period length was overcompensated with increasing temperature, suggesting that mechanisms controlling period lengthening may be linked to exaggerated temperature compensation (Fig. 4).

The circadian properties of *Wolffiella* appear to mimic *Arabidopsis* clock gene mutants. Similar to *W. hyalina*, the *prr7prr9* and *cca1lhy* double mutants show much longer periods at higher temperatures under constant light conditions; these double mutants show exaggerated temperature compensation of period length (Salomé *et al*., 2010; Shalit-Kaneh *et al*., 2018). With respect to the stability of circadian rhythms, the *gi* single mutant, *cca1lhy* double mutant, and *cca1lhyrve4rve6rve8* quint mutant show lower stability at higher and lower temperatures than at moderate temperatures (Gould *et al*., 2006; Shalit-Kaneh *et al*., 2018). The similarities in circadian properties between *Wolffiella* species and *Arabidopsis* loss-of-function mutants suggest that some of these clock genes may be malfunctioning in *Wolffiella* plants. Interestingly, *W, australiana* in the genus *Wolffia*, which is the closest relative to the *Wolffiella* genus, lacks several clock genes in its genome including some of PRRs (Michael *et al*., 2021). Thus, the lack of these clock-related genes may be involved in the instability of circadian rhythms and period overcompensation in *Wolffiella* plants.

### Relationship between circadian properties and geographical factors

*Lemna* plants showed more stable circadian rhythms than *Wolffiella* plants, with the former genus widely distributed in low- to high-latitude areas (Landolt, 1986). This wider distribution may be related to circadian rhythms with higher stability. The *Lemna* species found in warmer climates tended to show more unstable rhythms (Table S2); *L. valdiviana* inhabits tropical areas and showed the lowest stability among the *Lemna* species we observed; and *L. minuta*, the most closely related species to *L. valdiviana*, mainly inhabits higher latitudes, and its circadian rhythms were more stable than those of *L. valdiviana*. Together with the low stability of *Wolffiella* plants inhabiting low latitudes, the destabilization of circadian rhythms may be related to tropical and low-latitude environments. Such a relationship has also been discussed in marine cyanobacteria. *Prochlorococcus* inhabits low-latitude oceans while its closest marine relative *Synechococcus* inhabits a wide range of oceans up to high latitudes (Flombaum *et al*., 2013). Interestingly, *Synechococcus* maintains a basic set of clock genes while *Prochlorococcus* lacks an essential clock gene (*kaiA*) for circadian rhythmicity (Johnson & Egli, 2014). These findings suggest that the stability of circadian rhythms is important for inhabiting wide areas of the Earth. In addition to destabilization, the lengthening of plant circadian periods may be related to low-latitude environments. Indeed, very long period lengths (leaf movement) are observed in leguminous plants in tropical areas as well as in *W. hyalina* and *W. repanda* (Mayer, 1966). Thus, interspecific divergence in plant circadian rhythm stability appears to be related to species’ natural distributions. It should be noted that various species of *Lemna* and *Wolffiella* have overlapping distributions (Landolt, 1986). It would be interesting to determine whether differences in the stability of plant circadian rhythms are related to survival under a range of environmental conditions, such as in high-latitude areas.

### Morphological differences between *Lemna* and *Wolffiella* and their relation to the circadian properties

In *Arabidopsis*, it has been reported that different tissues have different circadian properties (Endo, 2016). In particular, the circadian clock of vascular tissue is robust and affects the circadian clocks of other tissues (Endo *et al*., 2014). In duckweed, *Lemna* and *Wolffiella* are morphologically distinct. Figure 6 shows the phylogenetic relationships between species from these two genera along with their morphological and circadian characteristics. Compared to *Lemna, Wolffiella* species have lost not only roots but also frond nerves, and they lack vascular tissue (Les *et al*., 1997). The observed unstable rhythms of *Wolffiella* plants under constant light conditions at high temperature appear to be linked to the degeneration of vascular tissue in this genus. Within the genus, *W. hyalina* and *W. repanda* have filament tracheids in the stamen while these are lacking in other *Wolffiella* species (Les *et al*., 1997). Interestingly, these two species exhibited relatively stable rhythms under constant conditions. Furthermore, *L. minuta* and *L. valdiviana*, which showed dampened rhythms under constant light conditions at high temperature, have also lost filament tracheids. Notably, no flowering was observed during the experiment period, which suggests that the loss of ability to differentiate filament tracheid may be linked to the destabilization of circadian rhythms in duckweed; genes involved in tracheid differentiation may influence the stability of circadian rhythms.

**Fig. 6.**
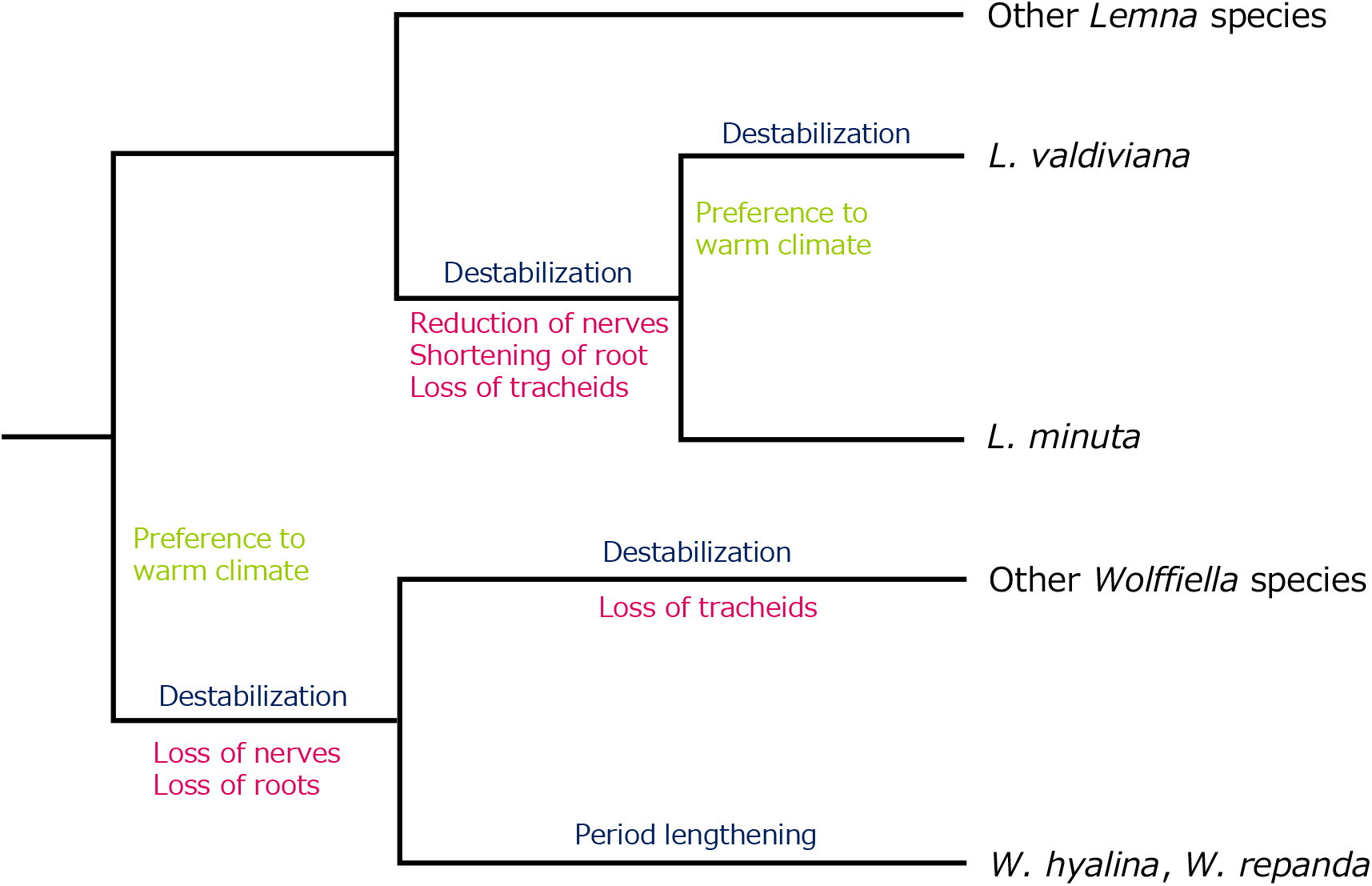
Schematic diagram representing an evolutionary scenario for the degeneration of circadian rhythms in duckweed. Evolutionary events of the circadian properties (blue), morphological features (red), and climate adaptation (green) are noted on a simplified phylogenetic tree.

Overall, our study reveals that interspecific divergence in the circadian properties of duckweed species primarily reflects phylogenetic relationships, and is likely to be related to geographical factors such as climate. Its evolutionary process will be approached through the comparative analysis of Lemnoideae and Wolffioideae genomes. It may be the case that genes controlling nerves or filament tracheids are also responsible for circadian stability. Furthermore, the fact that duckweed plants at low latitudes tended to show dampened/unstable rhythms at high temperatures raises the possibility that the degeneration of circadian rhythms generally occurs in plants in low-latitude areas. However, as yet, there are relatively few studies on the circadian rhythms of plants in the tropics (Mayer, 1966; Malpas & Jones, 2016). Further studies are, therefore, required to examine the circadian rhythms of plants in the tropics to contribute to our understanding of circadian adaptations in more stable environments.

## Acknowledgments

We thank Dr. Masaaki Morikawa, Dr. Walter Lämmler, and Dr. Eric Lam for providing us with duckweed plants. This work was supported in part by the Japan Society for the Promotion of Science KAKENHI (Grant numbers 19H03245, 21K18239), the Japan Science and Technology Agency (JST) ALCA and SATREPS to TO, and KAKENHI (Grant number 20K06342) to SI, and KAKENHI (Grant number JP21J15792) to MI.

## Author contributions

MI and TO planed and designed the research. MI performed experiments. MI analyzed the data. SI created a reporter construct. MI and TO wrote the manuscript.

## Supporting information

**Fig. S1.**
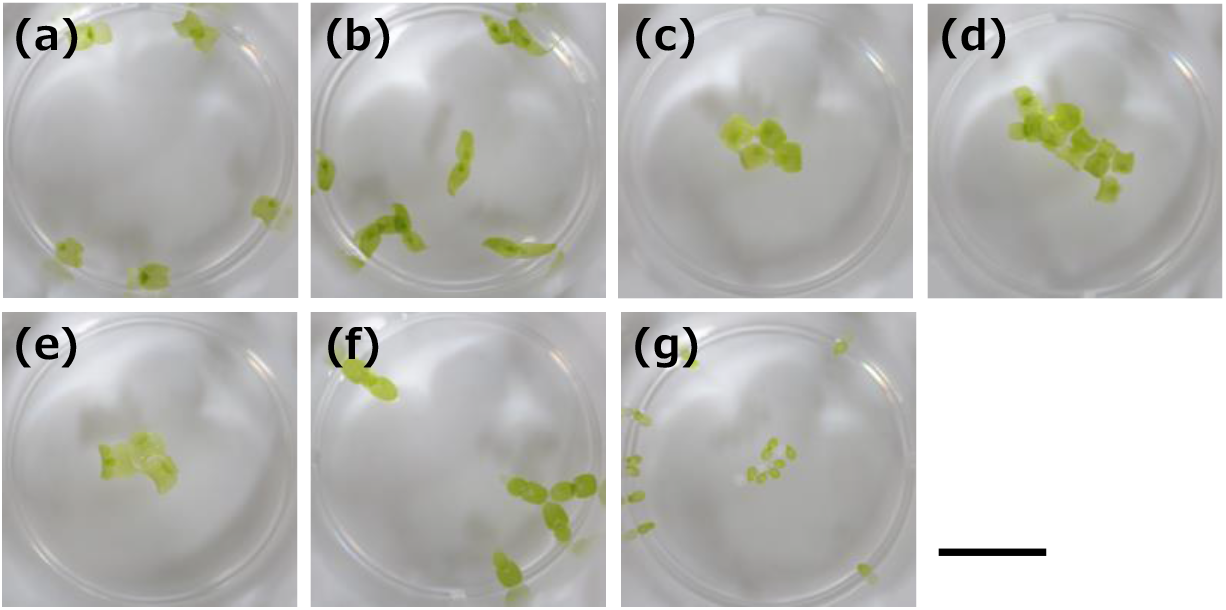
Morphological characteristics of the genus *Wolffiella*. (a) *W. lingulata*, (b) *W. oblonga*, (c) *W. caudata*, (d) *W. neotropica*, (e) *W. welwitschii*, (f) *W. hyalina*, and (g) *W. repanda*. Scale bar = 1 cm.

**Fig. S2.**
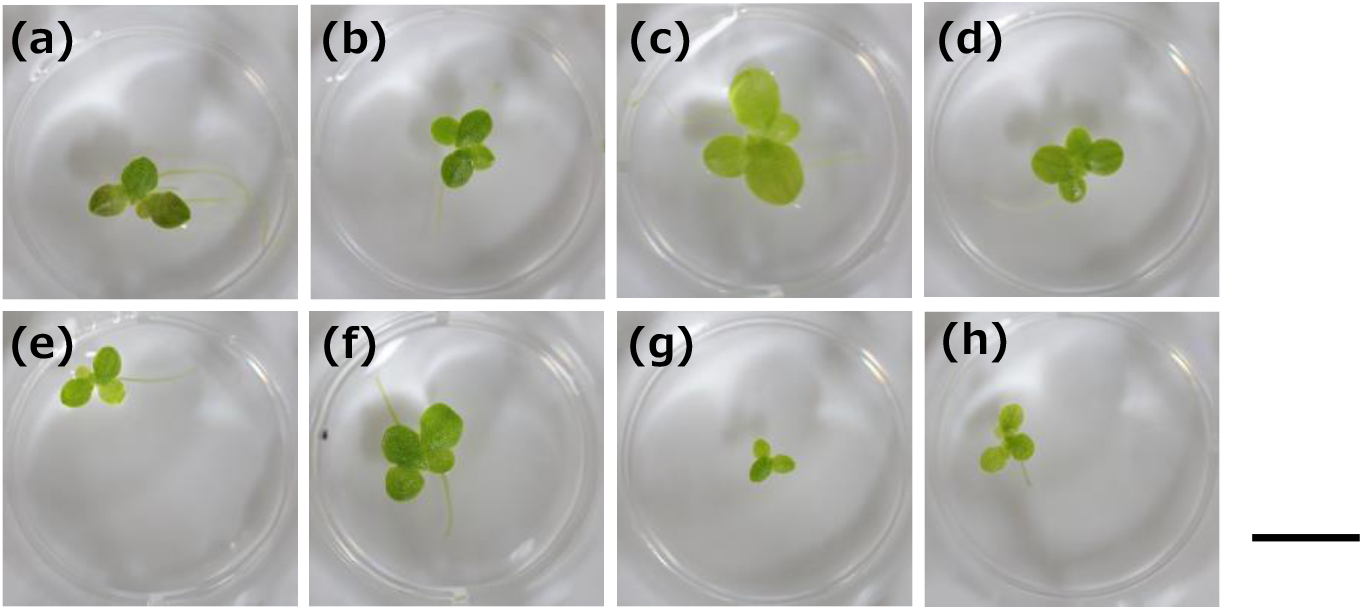
Morphological characteristics of the genus *Lemna*. (a) *L. obscura*, (b) *L. turionifera*, (c) *L. japonica*, (d) *L. minor*, (e) *L. disperma*, (f) *L. gibba*, (g) *L. valdiviana* and (h) *L. minuta*. Scale bar = 1 cm.

**Fig. S3.**
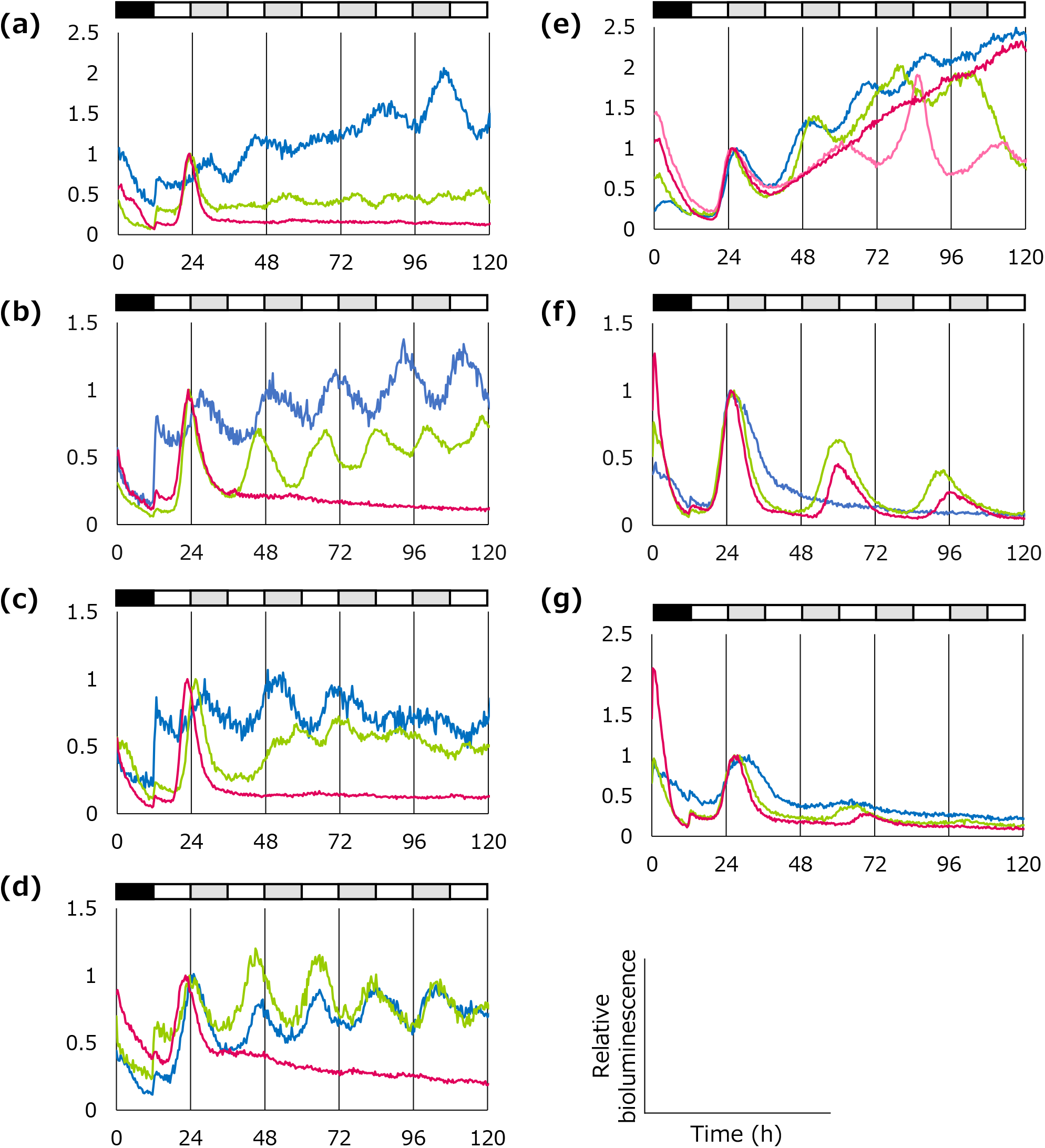
Bioluminescence of *AtCCR2::LUC+* under constant light conditions at various temperatures. *AtCCR2::LUC+* was introduced into (a) *W. lingulata*, (b) *W. oblonga*, (c) *W. caudata*, (d) *W. neotropica*, (e) *W. welwitschii*, (f) *W. hyalina*, and (g) *W. repanda*. Red, green, and blue colors indicate 30, 25, and 20 °C, respectively. Plants cultured in NF medium under constant light conditions were subjected to gene transfection and then entrained to two 12-h dark/12-h light cycles and released into constant light conditions; white and black bars indicate light and dark periods, respectively. The representative time-series data of two replicates in one experiment are shown for each species/condition. The data are presented as relative values with the highest value between 12 h and 36 h set to 1.

**Fig. S4.**
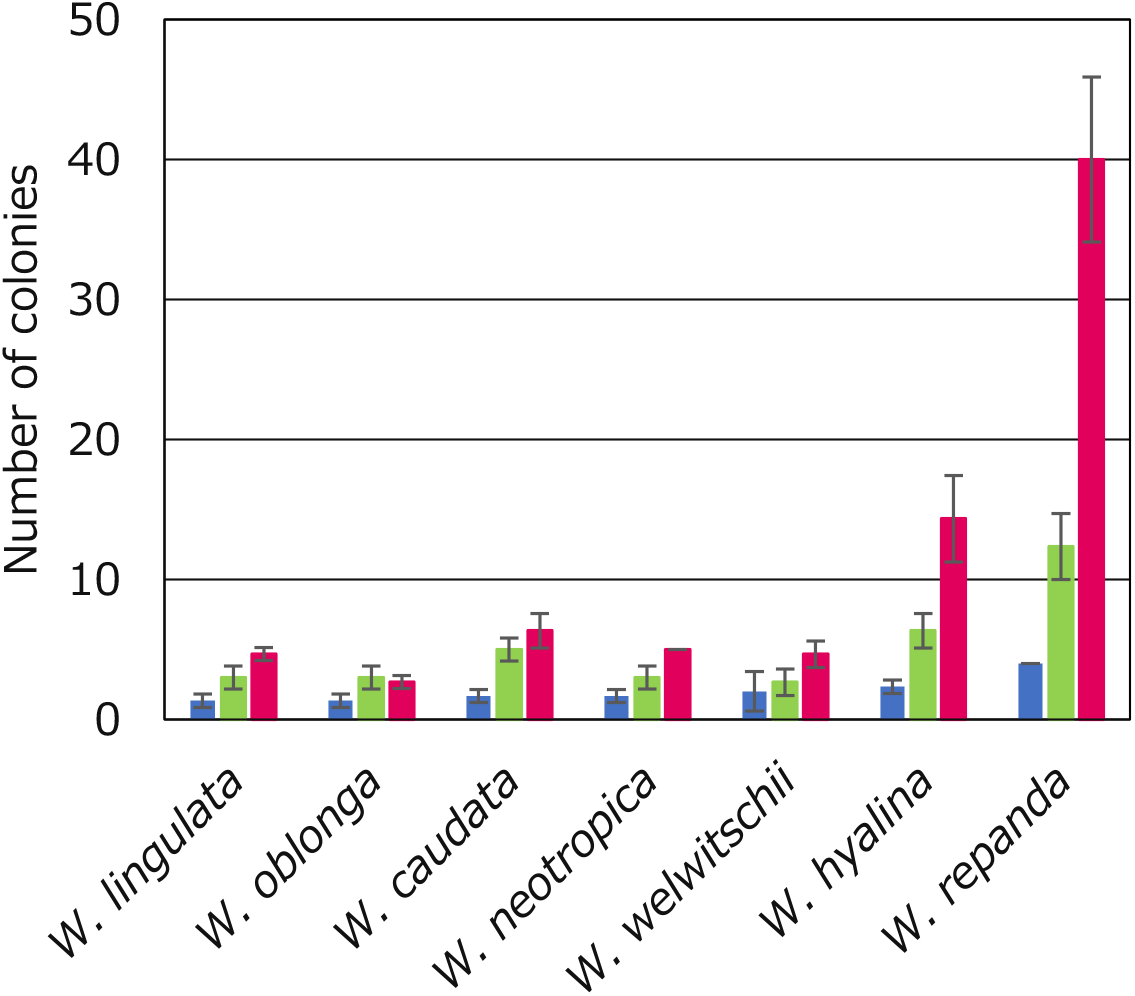
Comparison of growth at different temperatures. Plants were precultured in 12-h light/12-h dark conditions for two days at 20, 25, and 30 °C. After preculture, plants were placed in 12-well plates (one colony per well) and grown under constant light conditions at each temperature. After one week, the number of plant colonies in each well was counted. Data represent mean ± SD (*n* = 3). The red, green, and blue colors indicate 30, 25, and 20 °C, respectively.

**Fig. S5.**
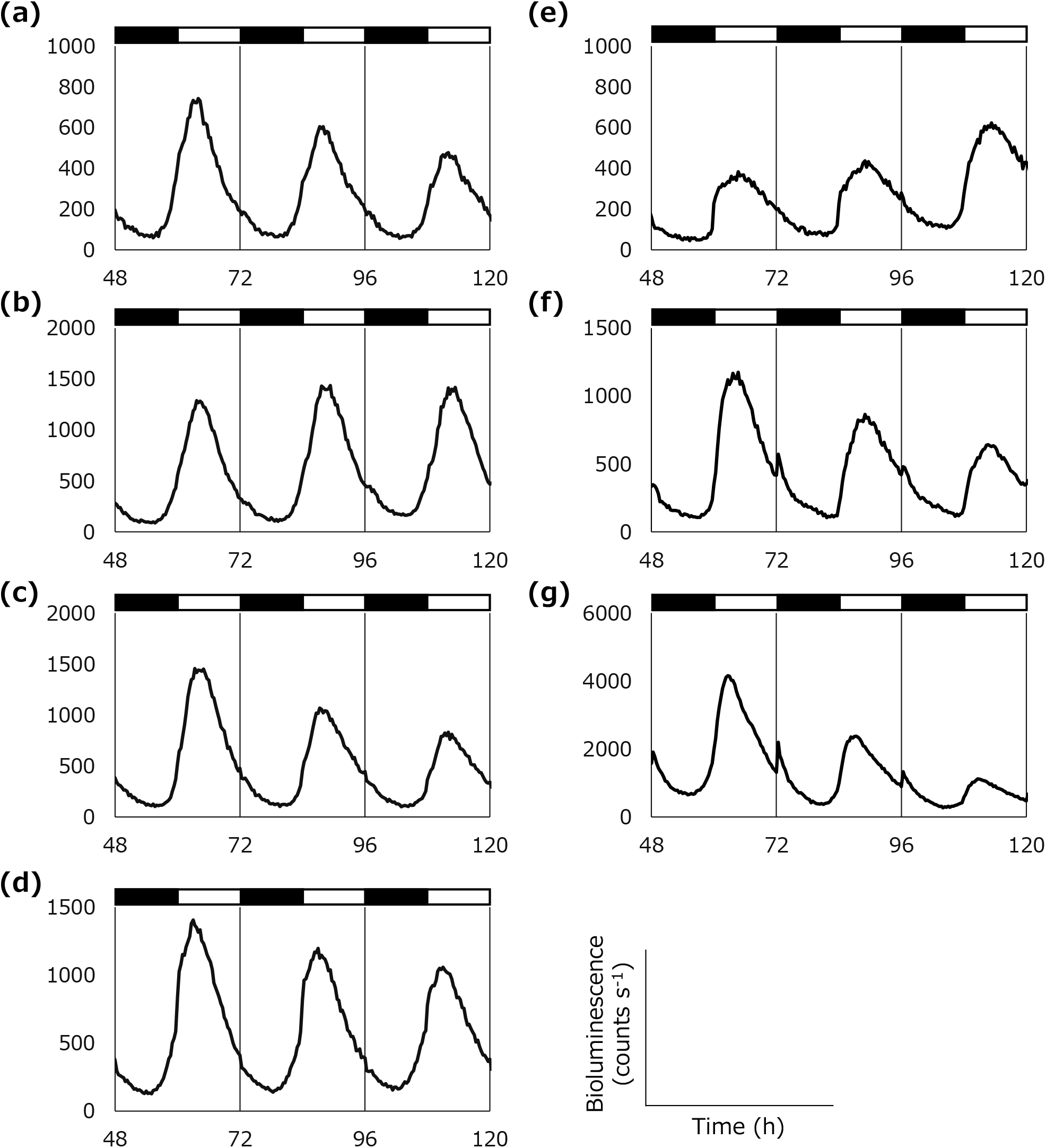
Bioluminescence of *AtCCA1::LUC+* under 12-h dark/12-h light conditions at 30 °C. *AtCCA1::LUC+* was introduced into (a) *W. lingulata*, (b) *W. oblonga*, (c) *W. caudata*, (d) *W. neotropica*, (e) *W. welwitschii*, (f) *W. hyalina*, and (g) *W. repanda*. Plants cultured in NF medium under constant light conditions were subjected to gene transfection, and bioluminescence rhythms were measured under 12-h dark/12-h light conditions at 30 °C. White and black bars indicate light and dark periods, respectively. The representative time-series data of three replicates in one experiment are shown for each species/condition.

**Table S1.** Circadian rhythm stability of duckweed plants under each experimental condition.

| Species | 20°C |  | 25°C |  | 30°C |  |
| --- | --- | --- | --- | --- | --- | --- |
|  | LL | DD | LL | DD | LL | DD |
| *W. lingulata* | R | D | D | R | AR | R |
| *W. oblonga* | R | D | R | R | D | R |
| *W. caudata* | R | D | D | R | AR | R |
| *W. neotropica* | D | D | R | R | AR | R |
| *W. welwitschii* | D | D | R | R | AR | R |
| *W. hyalina* | D | D | R | R | R | R |
| *W. repanda* | D | D | R | R | D | R |
| *L. obscura* | D | R | R | R | R | R |
| *L. turionifera* | D | R | R | D | R | R |
| *L. japonica* | D | R | R | R | R | R |
| *L. minor* | D | R | R | R | R | R |
| *L. disperma* | R | R | R | R | R | R |
| *L. gibba* | R | R | R | R | R | R |
| *L. valdiviana* | D | D | R | R | D | D |
| *L. minuta* | R | D | R | R | D | R |

AR: arrhythmic, D: dampened rhythm, R: rhythmic

**Table S2.** Correspondence between the Köppen climate classification of duckweed plants and circadian rhythm stability under each experimental condition.

| Species | Köppen<br>climate<br>classification | 20°C |  | 25°C |  | 30°C |  |
| --- | --- | --- | --- | --- | --- | --- | --- |
|  |  | LL | DD | LL | DD | LL | DD |
| *W. neotropica* | Aw | D | D | R | R | AR | R |
| *W. caudata* | Am | R | D | D | R | AR | R |
| *L. valdiviana* | Am | D | D | R | R | D | D |
| *W. hyalina* | BS | D | D | R | R | R | R |
| *W. repanda* | BS | D | D | R | R | D | R |
| *W. welwitschii* | BW | D | D | R | R | AR | R |
| *W. oblonga* | Cfa | R | D | R | R | D | R |
| *L. obscura* | Cfa | D | R | R | R | R | R |
| *L. japonica* | Cfa | D | R | R | R | R | R |
| *L. minuta* | Cfb | R | D | R | R | D | R |
| *W. lingulata* | Cs | R | D | D | R | AR | R |
| *L. turionifera* | Cs | D | R | R | D | R | R |
| *L. disperma* | Cs | R | R | R | R | R | R |
| *L. gibba* | Cs | R | R | R | R | R | R |
| *L. minor* | Df | D | R | R | R | R | R |

AR: arrhythmic, D: dampened rhythm, R: rhythmic

